# BMP antagonism is required for mandible outgrowth in zebrafish

**DOI:** 10.64898/2026.03.11.711234

**Authors:** Hung-Jhen Chen, Jennifer Dukov, Trevor Llyod, Pengfei Xu, D’Juan T. Farmer

**Affiliations:** Department of Molecular, Cell and Developmental Biology, University of California, Los Angeles; Department of Orthopaedic Surgery, University of California, Los Angeles; Department of Orofacial Sciences and Program in Craniofacial Biology, University of California, San Francisco, San Francisco, CA, USA; Howard Hughes Medical Institute

**Keywords:** zebrafish, Meckel’s cartilage, mandible, BMP signaling, chondrocyte differentiation

## Abstract

The Meckel’s cartilage (MC) is a fundamental component of mandibular development across vertebrates. In mammals, MC is transient and functions primarily as an early template for mandibular ossification, whereas other vertebrates, including zebrafish, retain MC within the mandible throughout life. Despite its importance, the requirements for MC in sustaining mandibular growth and how signaling pathways implicated in MC development contribute to this process remain unclear. Here, we investigated the dosage-dependent roles of BMP antagonists during zebrafish MC development using mutant alleles of *grem1a*, *nog2*, and *nog3*. Compound mutant adults exhibited fully penetrant mandibular truncation. MC shortening emerged after early larval stages, indicating a requirement for BMP antagonism to sustain cartilage growth. Chondrocyte number remained unchanged as phenotypes developed, but mutants displayed disorganized cartilage morphology and increased chondrocyte volume. Molecular analyses revealed reduced *col2a1a* domains and expanded *ihha* and *col10a1a* expression, consistent with ectopic hypertrophic-like differentiation. Constitutive activation of BMP receptor signaling in chondrocytes recapitulated these phenotypes. Although osteogenesis appeared unaffected by 14 dpf, loss of a *tnn*⁺ skeletal mesenchyme population was observed. Together, these findings demonstrate that BMP antagonists sustain MC growth by regulating chondrocyte differentiation and cartilage organization to support mandibular growth in non-mammalian vertebrates.

**Summary Statement:** This study leverages zebrafish to define the cellular and molecular mechanisms by which BMP antagonism sustains mandibular growth.

## Introduction

The mandible is required for proper feeding and communication and disruption of its development leads to congenital anomalies in humans. It arises from neural crest cells that populate the mandibular pharyngeal arch (Chai et al., 2000). One of the earliest skeletal elements to differentiate within this arch is Meckel’s cartilage (MC), a structure that acts as a template for mandibular development. In mammals, MC exhibits regionally distinct fates along the proximal–distal axis (Anthwal et al., 2013; Svandova et al., 2020). The proximal domain contributes to the malleus and incus of the middle ear, the intermediate portion degenerates without forming bone, and the distal region undergoes endochondral ossification. By early postnatal stages, remnants of MC have disappeared. In contrast, in other vertebrates like the zebrafish, MC persists into adulthood and remains an integral component of the mandible (Anthwal et al., 2013; DeLaurier, 2019; Svandova et al., 2020), although its functional requirements at later stages of development remain untested. Despite these species-specific differences, zebrafish MC has served as a valuable model for identifying conserved mechanisms governing cartilage formation and growth.

MC serves as an essential template for intramembranous ossification. Perturbations affecting neural crest migration, mandibular arch patterning, or early skeletal differentiation can disrupt mandibular growth and result in micrognathia, a congenital condition characterized by reduced jaw size (Adel Al-Lami et al., 2016; Bonatto Paese et al., 2021; Chen et al., 2019; Iwaya et al., 2023; Parada et al., 2015; Stottmann et al., 2001). Genetic studies in mice demonstrate that deletion of the essential chondrogenic regulator *Sox9* produces a reduced mandible, suggesting that although MC is dispensable for intramembranous ossification, it is critical for determining mandibular size (Mori-Akiyama et al., 2003). Defining how MC growth is regulated therefore remains a fundamental question in craniofacial biology.

BMP signaling is a key regulator of skeletal patterning and growth, and modulates proliferation, cell survival, and differentiation. Deletion of *Bmpr1a* and *Bmpr1b* in chondrocytes results in widespread impairment of early chondrogenesis (Yoon et al., 2005). Within the craniofacial complex, BMP4 produced by pharyngeal epithelia promotes survival of mandibular mesenchyme and supports mandibular outgrowth (Liu et al., 2005). Neural crest-derived BMP2 and BMP4 further influence mandibular outgrowth, and elevated BMP activity disrupts craniofacial morphogenesis (Bonilla-Claudio et al., 2012). Experimental manipulations in avian and mammalian systems reveal stage-specific effects of BMP signaling on cartilage differentiation and osteoblast maturation (Hu et al., 2008; Merrill et al., 2008), underscoring the requirement for tightly regulated BMP activity during mandibular development.

BMP antagonists, which include members of the Gremlin, Noggin, Chordin, and Follistatins, refine pathway activity by binding BMP ligands and restricting receptor activation. Loss of the antagonist Noggin results in expansion of MC and consequently, mandibular thickening (Matsui and Klingensmith, 2014; Stottmann et al., 2001; Wang et al., 2013), while increased BMP receptor signaling phenocopies these effects (Wang et al., 2013). Combined disruption of Noggin and Chordin produces mandibular hypoplasia and variable micrognathia, contributed to a role for balanced BMP signaling for neural crest survival and early cartilage development (Matsui and Klingensmith, 2014; Stottmann et al., 2001). However, if and how BMP signaling is required to ensure sustained MC growth is unclear and whether BMP antagonists’ function cooperatively to support MC development remain poorly defined.

Here, we employ the zebrafish model to investigate dosage-dependent requirements of BMP antagonists during MC development. Leveraging duplicated antagonist genes, we analyze mutants for *nog2*, *nog3*, and *grem1a*. We show that adult *grem1a*^-/-^; *nog2*^-/-^; *nog3*^+/-^ mutants exhibit pronounced mandibular truncation, contrasting with phenotypes observed in mice. Developmental analyses reveal overlapping antagonist expression within MC chondrocytes and its surrounding skeletal mesenchyme. While early larval skeletal size is preserved, MC growth becomes significantly reduced by 14 dpf, indicating a requirement for BMP antagonism to sustain cartilage growth during post-patterning stages. Mutants display disrupted chondrocyte organization and altered chondrocyte differentiation associated with increased cell size and the acquisition of a hypertrophic-like state, without impairing early intramembranous ossification. Truncations emerged earlier in *grem1a*^-/-^; *nog2*^-/-^; *nog3*^-/-^ mutants, which fail to survive to later stages, reinforcing the sensitivity of MC to BMP signaling. Finally, constitutive activation of *Bmpr1a* in chondrocytes is sufficient to induce hypertrophic programs, supporting a conserved role for balanced BMP signaling in regulating chondrocyte differentiation. Together, these findings demonstrate that BMP antagonists exert dosage-sensitive control over MC growth and differentiation, and reveal a prolonged consequence of disrupting BMP signaling on mandible outgrowth. These results provide new insight into how antagonistic regulation of BMP signaling ensures proper craniofacial growth and size.

## Results

### Mutations in BMP antagonists cause mandible truncations

Previously, we identified co-expression of the BMP antagonists *grem1a*, *nog2*, and *nog3* during cranial suture formation and demonstrated that their activation is required for timely suture development (Farmer et al., 2024). We next asked if gremlins and noggins have additional functions during craniofacial development. When examining *grem1a*^-/-^; *nog2*^-/-^; *nog3*^+/-^ zebrafish, which are adult viable unlike their *grem1a*^-/-^; *nog2*^-/-^; *nog3*^-/-^ siblings, for gross craniofacial abnormalities, we observed several defects in the craniofacial skeleton. These included delayed maturation of the opercular bones despite animals being size-matched. Most prominently, mutants exhibited a shortened yet mobile mandible, which we investigated further in this study (**Fig. 1A, Movie 1**). MicroCT analysis confirmed the presence of an unfused jaw joint accompanied by marked mandibular truncation (**Fig. 1A**). Skeletal staining with Alizarin Red (bone) and Alcian Blue (cartilage) further verified that mutant fish exhibited a truncated and widened mandible, while cartilage at the jaw joint remained intact (**Fig. 1B**). Quantification of lower jaw length relative to the upper jaw revealed that *grem1a*^-/-^; *nog2*^+/-^; *nog3*^+/-^ zebrafish (termed controls) displayed mandibular extension beyond the upper jaw by approximately 150 µm. In contrast, all *grem1a*^-/-^; *nog2*^-/-^; *nog3*^+/-^ zebrafish (termed dhomo; het) showed pronounced reductions, with the lower jaw positioned on average 520 µm behind the upper jaw, reflected as negative values (**Fig. 1C**). Together, these findings demonstrate that BMP antagonists are required to achieve proper mandibular size.

**Figure 1.**
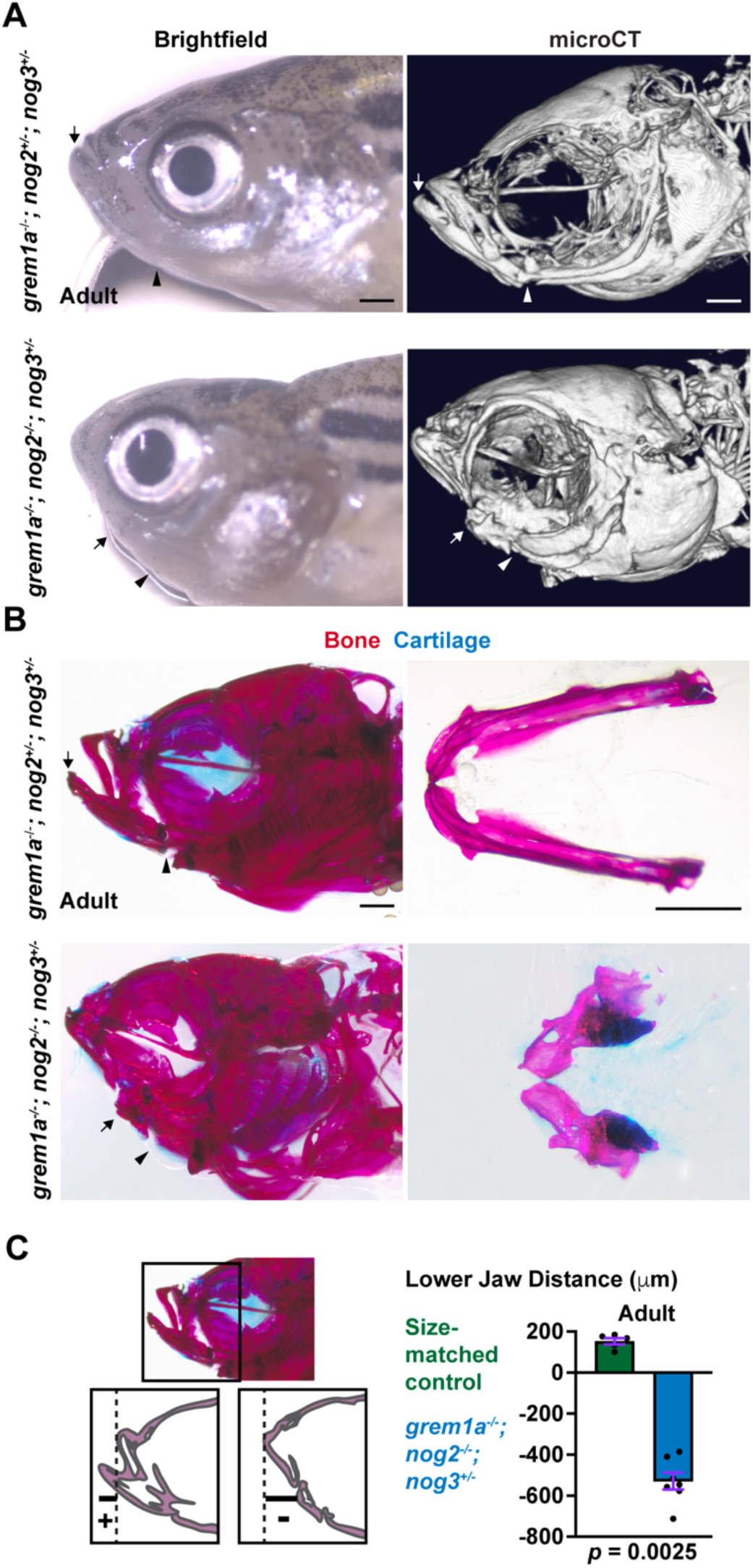
Compound mutants for BMP antagonists have a truncated mandible. **(A)** Live image (n = 6 per genotype) and microCT (n = 2 per genotype) of *grem1a*^-/-^; *nog2*^+/-^; *nog3*^+/-^ and *grem1a*^-/-^; *nog2*^-/-^; *nog3*^+/-^ adults show reduced mandibles. **(B)** Skeletal staining of bone (red) and cartilage (blue) (n = 4 per genotype) shows adult *grem1a*^-/-^; *nog2*^-/-^; *nog3*^+/-^ mutants display truncated lower jaw compared to size-matched controls. Dissected lower jaws are shown in the right panel and highlight the malformed mutant lower jaw compared to size-matched control. Jaw joint cartilage persists in the mutant lower jaw and processes of the dentary bone are still visible in the mutants. **(C)** Quantification of the horizontal distance between the anterior tip of the lower jaw to the anterior tip of the upper jaw with positive values indicating the lower jaw extension beyond the upper jaw and negative values indicating the lower jaw truncation behind the upper jaw (n =5 controls and n = 7 mutants) Arrows point to the anterior of the lower jaw and arrowheads indicated the posterior of the lower jaw. Scale bars, 500 µm. Error bars represent SEM.

### BMP antagonists are expressed with MC chondrocytes and skeletal mesenchyme

To investigate how BMP antagonists contribute to mandibular size, we analyzed published single-cell datasets of zebrafish neural crest derivatives across developmental stages (Fabian et al., 2022). At 2 dpf, *grem1a* expression was not detected, whereas *nog2* was broadly expressed within *postnb⁺* mesenchyme, with limited co-expression of *nog3* (**Fig. S1A**). Markers of differentiated osteoblasts (*ifitm5*) and chondrocytes (*mia*) were not yet observed. By 5 dpf, *nog2* and *nog3* were co-detected within *mia⁺* chondrocytes, while *grem1a*, *nog2*, and *nog3* were co-expressed within *postnb⁺* mesenchyme. These expression patterns persisted at 14 dpf and 60 dpf, suggesting sustained roles for BMP antagonists during craniofacial development. To validate these transcriptional profiles, we performed combinatorial RNAscope in situ hybridization. At 6 dpf, *nog2* and *nog3* transcripts were detected within MC chondrocytes, whereas *grem1a* expression was comparatively low (**Fig. 2A, C**). Notably, *grem1a* and *nog3* were enriched in chondrocytes at the jaw joint. All three antagonists were also detected in the skeletal mesenchyme along the lateral aspects of MC. Imaging of the entire lower jaw at this stage demonstrated widespread *nog2* expression across larval cartilages. *nog3* showed lower overall chondrocyte expression but was concentrated at articulating cartilage regions, whereas *grem1a* was selectively enriched in the outer MC mesenchyme (**Fig. S1B**). At 14 dpf, these expression domains were maintained, with detection across MC and a pronounced increase in *grem1a* expression within the lateral mesenchyme (**Fig. 2B, C**). Together, these findings demonstrate overlapping and sustained expression of BMP antagonists within MC chondrocytes and surrounding mesenchyme, consistent with roles in regulating MC growth.

**Figure 2.**
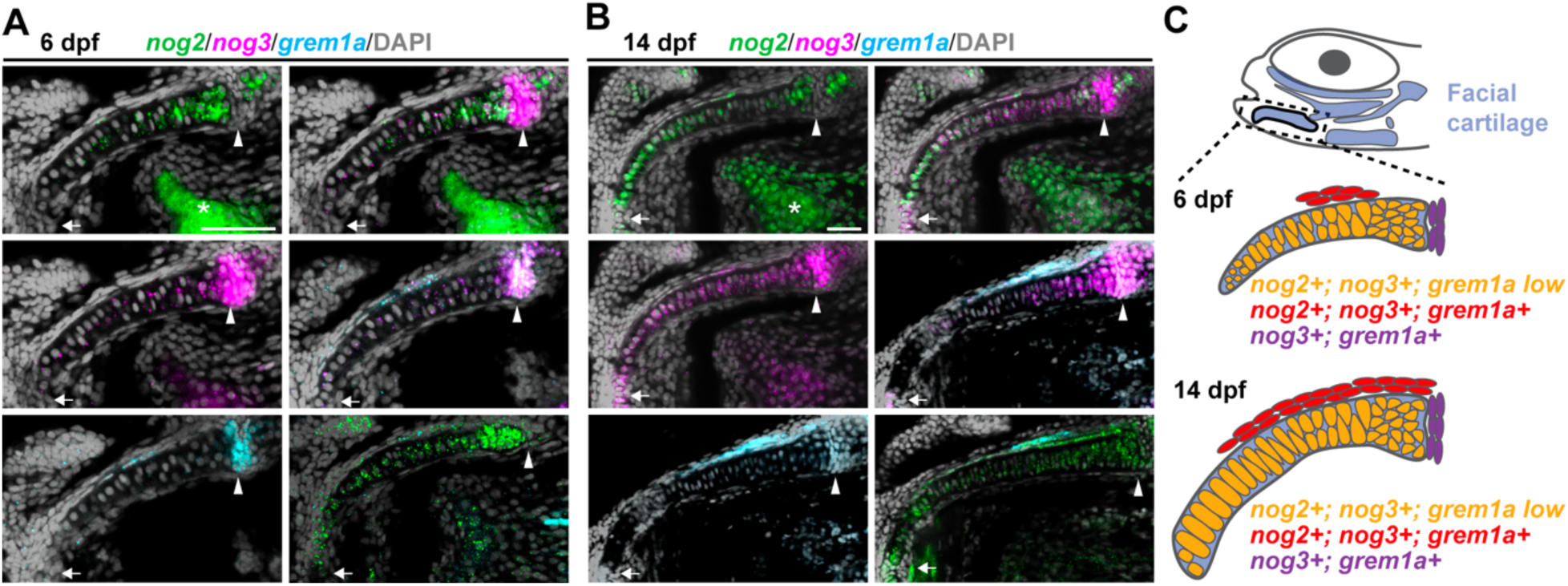
Noggin and Gremlin genes are co-expressed in chondrocytes and adjacent skeletal mesenchyme. (**A, B**) Single Z-plane confocal images of Meckel’s cartilage following in situ hybridization for *nog2* (green), *nog3* (magenta), and *grem1a* (blue), counter-stained with DAPI (gray), at 6 dpf (A) and 14 dpf (B). BMP antagonist transcripts are detected in chondrocytes as well as in a discrete population of skeletal mesenchyme immediately adjacent to Meckel’s cartilage. (**C**) Schematic summary of expression profiles of BMP antagonists in and around Meckel’s cartilage. Arrows point to the anterior of the Meckel’s cartilage and arrowheads indicated the posterior of the Meckel’s cartilage. All stains were completed in biological triplicates. Asterisk marks autofluorescence lacking punctate RNAScope signal. Scale bars, 50 µm.

### MC truncations can arise after proper establishment of the larval skeleton

To determine when mandibular truncation arises, we examined skeletal development at 6 dpf and 14 dpf using Alcian Blue and Alizarin Red staining in *grem1a*^-/-^; *nog*^+/-^; *nog3*^+/-^ (control), *grem1a*^-/-^; *nog2*^-/-^; *nog3*^+/-^ (dhomo; het), and *grem1a*^-/-^; *nog2*^-/-^; *nog3*^-/-^ (triple homo) zebrafish. As triple homo mutants die during larval stages, we analyzed them only at 6 dpf. At 6 dpf, the size and morphology of the larval skeleton were indistinguishable between control and dhomo; het zebrafish, while MC in triple homo mutants appeared shorter and thicker (**Fig. 3A**). By contrast, at 14 dpf, dhomo; het MCs were visibly shortened and no longer protruded beyond the neurocranium (**Fig. 3B**). Flat-mount preparations of control and dhomo; het mutant skeletons confirmed a selective truncation of MC, while other craniofacial elements appeared comparatively normal (**Fig. 3C**). Quantitative analysis showed no significant difference in MC extension past the neurocranium at 6 dpf between control and dhomo; het mutants but a significant reduction in MC extension in triple homo mutants (**Fig. 3D**). However, by 14 dpf, dhomo; het mutants exhibited a significant reduction in the extent to which MC extended past the neurocranium (**Fig. 3D**). Triple homo mutants showed a significant decrease MC size at 6 dpf. Together, these findings indicate that mandibular truncation can emerge progressively in response to reduced BMP antagonism in the larval skeleton and, at least in dhomo; het mutants, reflects impaired MC growth rather than early patterning defects.

**Figure 3.**
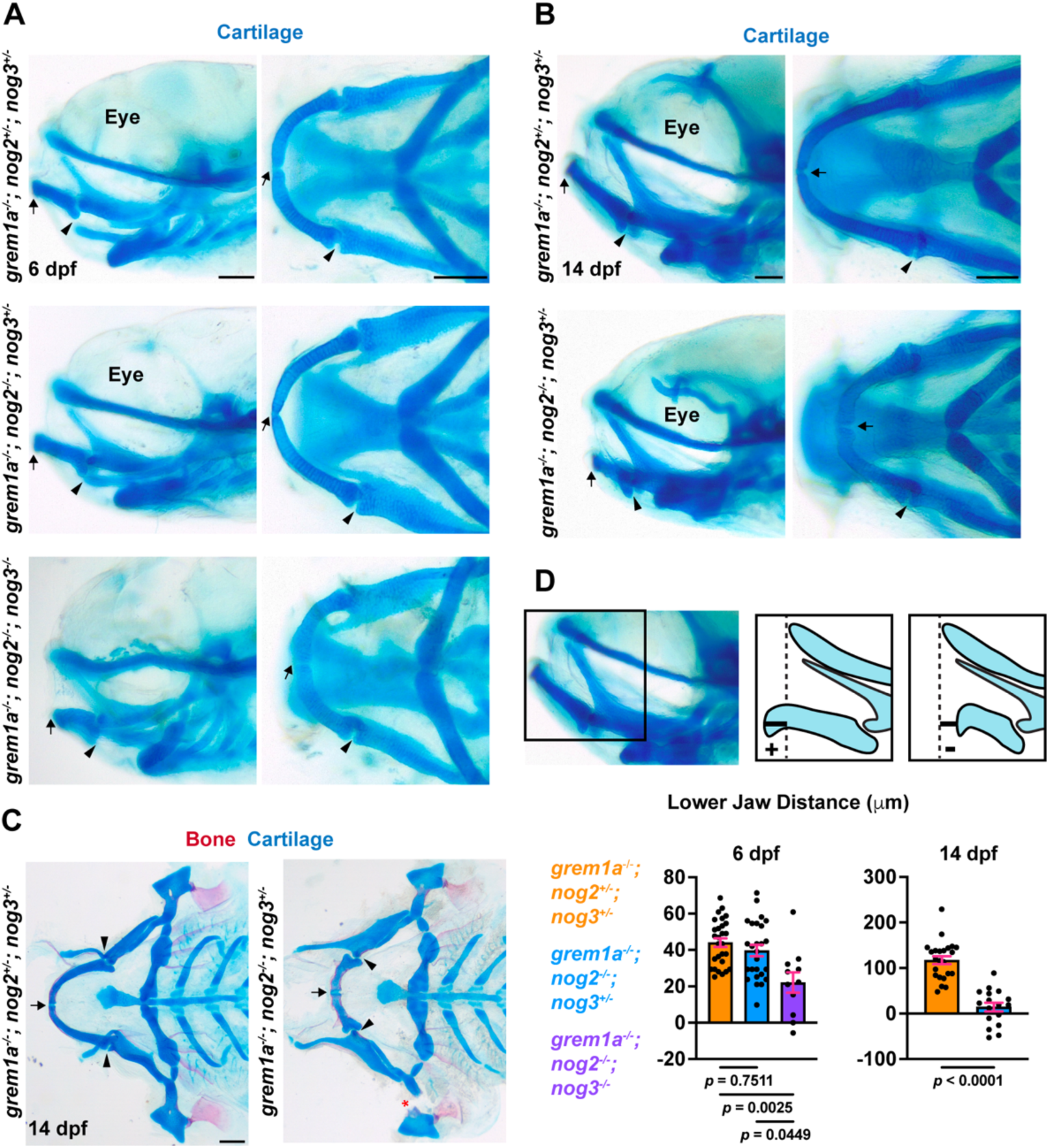
Lower jaw truncations in *grem1a; nog2; nog3* compound mutants emerge after the establishment of the larval skeleton. (**A**) Lateral (left) and ventral (right) views of Alcian blue stained larval skeletons show comparable skeletal morphology between *grem1a*^-/-^; *nog2*^+/-^; *nog3*^+/-^ and *grem1a*^-/-^; *nog2*^-/-^; *nog3*^+/-^ but reductions in *grem1a*^-/-^; *nog2*^-/-^; *nog3*^-/-^ zebrafish at 6 dpf (n = 28 control and 26 dhomo; het and 11 triple homo fish). (**B**) Lateral (left) and ventral (right) views of Alcian blue stained skeletons at 14 dpf reveal shortening of Meckel’s cartilage in *grem1a*^- /-^; *nog2*^-/-^; *nog3*^+/-^ zebrafish (n = 23 control and 17 dhomo; het fish). (**C**) Flat-mount of Alcian blue and Alizarin Red stained control and mutant skeletons demonstrate selective reduction in the Meckel’s cartilage size within the craniofacial skeleton (n = 6 per genotype). (**D**) Quantification of the horizontal distance between the anterior tip of Meckel’s cartilage and the anterior tip of the neurocranium. Positive values indicate that Meckel’s cartilage extends beyond the neurocranium, whereas negative values indicate that it falls posterior to the neurocranium. Distances are comparable at 6 dpf (n = 28 control and 26 dhomo; het fish) between *grem1a*^-/-^; *nog2*^+/-^; *nog3*^+/-^and *grem1a*^-/-^; *nog2*^-/-^; *nog3*^+/-^ fish, but reduced in *grem1a*^-/-^; *nog2*^-/-^; *nog3*^-/-^ fish (n = 11). By 14 dpf, the mutant lower jaw no longer extends as far anteriorly as controls (n = 23 control and 17 dhomo; het fish). Arrows point to the anterior of the Meckel’s cartilage and arrowheads indicated the posterior of the Meckel’s cartilage. Scale bars, 100 µm. Error bars represent SEM.

We next examined 3D morphometric features of MC. To visualize chondrocytes, control and mutant zebrafish were stained with Wheat Germ Agglutinin (WGA), which labels the chondrocyte extracellular matrix (Schlombs et al., 2003). Imaris-based 3D reconstructions were then used to quantify total MC volume (**Fig. 4A, B**). At 6 dpf, both dhomo; het and triple homo mutants exhibited a significant increase in MC volume relative to controls, with a slight but insignificant increase in triple homo mutants compared to dhomo; het mutants. By contrast, no obvious difference in MC volume was detected at 14 dpf between control and dhomo; het animals, despite clear reductions in MC length. In triple homo mutants at 6 dpf and dhomo; het mutants at 14 dpf, the MC exhibited an abnormal, uneven morphology consistent with disrupted cartilage architecture (**Fig. 4B**). To determine whether altered cartilage architecture contributed to this phenotype, we measured MC thickness along the dorsal-ventral and lateral-medial axes at 14 dpf (**Fig. 4C**). Compared to controls, dhomo; het mutant MCs were significantly thickened along both dimensions, suggesting that changes in cartilage organization accompany MC shortening. To assess whether cellular alterations underlie these morphological changes, we quantified chondrocyte volume within the intermediate MC domain (**Fig. 4D**). At both 6 and 14 dpf, mutant chondrocytes displayed significantly increased cell volume. This increase likely explains the elevated MC volume observed at 6 dpf and suggests altered chondrocyte differentiation in mutant MCs.

**Figure 4.**
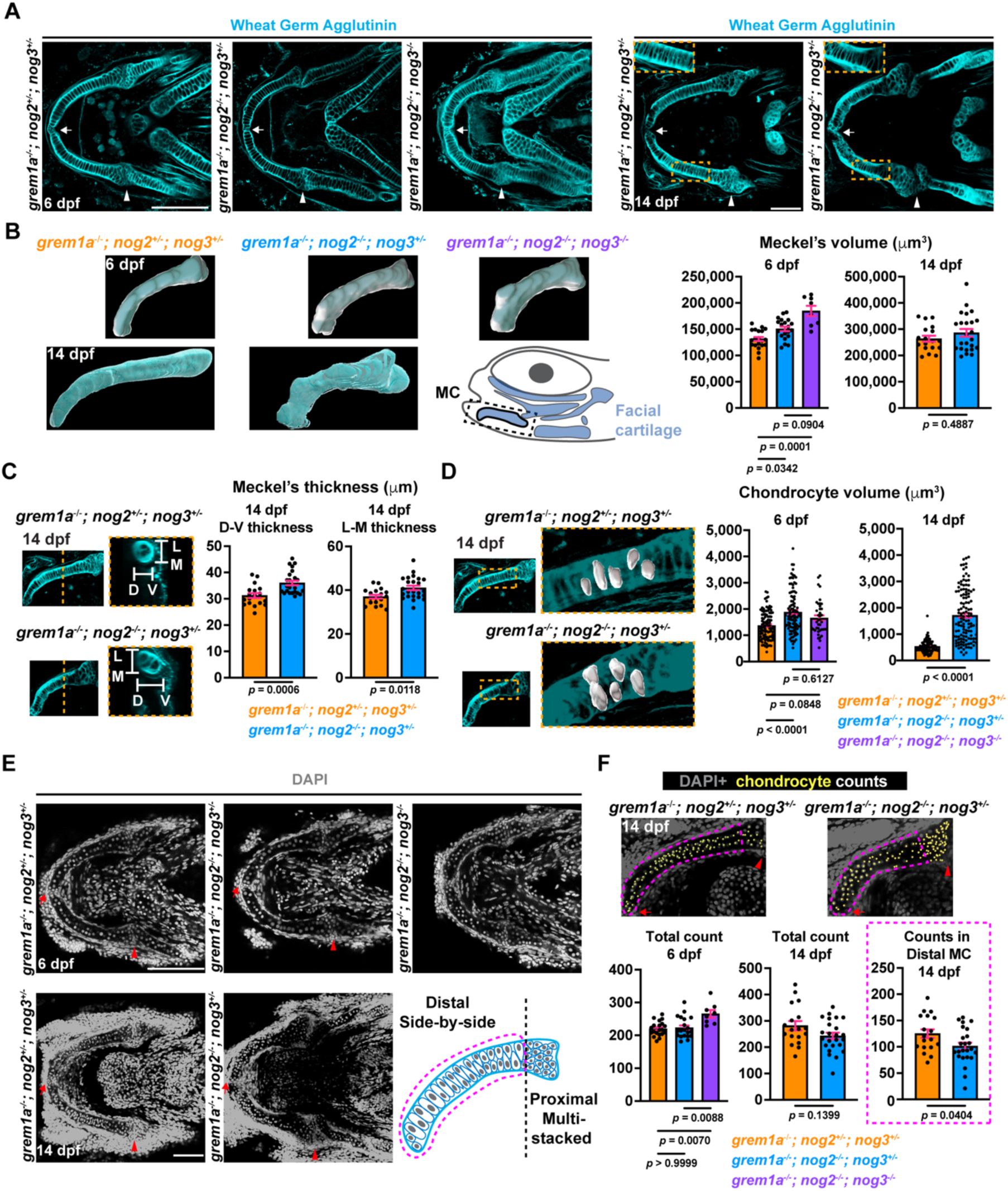
Altered cartilage morphology and increased chondrocyte volume underlie Meckel’s cartilage truncation in mutant zebrafish. (**A**) Single Z-plane confocal images of Wheat Germ Agglutinin (WGA) stained skeletons at 6 dpf and 14 dpf highlighting Meckel’s cartilage. Yellow dashed boxes indicate the regions shown in higher-magnification insets. Chondrocytes in mutants appear enlarged and less well aligned compared to controls. (**B**) Representative 3D reconstructions of 6 dpf and 14 dpf control and mutant Meckel’s cartilage generated from WGA staining to assess cartilage volume. Quantification of cartilage volume at 6 dpf (n = 20 control, 20 dhomo; het, and 8 triple homo Meckel’s cartilages), and 14 dpf (n = 18 control and 24 dhomo; het Meckel’s cartilages) is shown. Cartilage volume is increased at 6 dpf but comparable by 14 dpf, despite the truncated and irregular morphology of mutant Meckel’s cartilages. (**C**) Representative single Z-plane confocal images of WGA-stained Meckel’s cartilage at 14 dpf, with cross-sectional views illustrating cartilage thickness along the dorsal–ventral and lateral–medial axes. Quantification of thickness along each axis (n = 18 control and 24 dhomo; het Meckel’s cartilages) reveals increased thickness in mutants across both dimensions. (**D**) Representative 3D reconstructions of five chondrocytes from WGA-stained Meckel’s cartilage in control and mutant zebrafish at 14 dpf. Quantification of chondrocyte volume at 6 dpf (five cells per cartilage; n = 20 control, 20 dhomo; het, and 8 triple homo Meckel’s cartilages), and 14 dpf (five cells per cartilage; n = 18 control and 24 dhomo; het Meckel’s cartilages) shows increased cell size in mutants by 6 dpf that persists through 14 dpf. (**E**) Representative images of single Z-plane confocal images of DAPI stained control and mutant zebrafish at 6 and 14 dpf. (**F**) Quantification of DAPI+ chondrocyte numbers across full confocal z-stacks of Meckel’s cartilage (single Z-plane shown for representation) indicates that total chondrocyte number is comparable between control and dhomo; het larvae at 6 dpf (n = 20 control and 20 dhomo; het cartilages) and 14 dpf (n = 18 control and 24 dhomo; het cartilages). However, triple homo (n = 8 cartilages) displayed increased chondrocyte numbers at 6 dpf. When 14 dpf analyses are restricted to distal regions of Meckel’s cartilage where chondrocytes adopt a side-by-side organization, dhomo; het mutants exhibit reduced chondrocyte numbers relative to controls. Arrows point to the anterior of the Meckel’s cartilage and arrowheads indicated the posterior of the Meckel’s cartilage. Scale bars, 100 µm. Error bars represent SEM.

In dhomo; het fish, we detected an increase in mutant chondrocyte volumes despite no difference in total MC volume and truncated MC length at 14 dpf. To assess whether altered cell number contributes to MC truncation in mutant zebrafish, DAPI-stained control and mutant zebrafish were analyzed using ImageJ at 6 dpf or Imaris Spot detection at 14 dpf (**Fig. 4E, F**). Total chondrocyte number was unchanged at both 6 dpf and 14 dpf between control and dhomo; het fish (**Fig. 4F**). In contrast, total chondrocyte number was increased in triple homo fish, despite early signs of MC truncation. While total cell numbers were unchanged between control and dhomo;het fish, we observed an altered organization of chondrocytes in dhomo;het mutants, where chondrocyte volume was increased (**Fig. 4D**). Quantification of chondrocytes at 14 dpf where cellular organization transitions from a multi-stacked arrangement near the jaw joint to a side-by-side configuration revealed a significant decrease in chondrocyte number (**Fig. 4F**). These findings are consistent with impaired maintenance of chondrocyte expansion within MC.

In zebrafish, interstitial growth has been proposed as a mechanism contributing to Meckel’s cartilage elongation, particularly after initial establishment of MC and during later larval growth stages (Eames et al., 2013). To determine whether altered proliferation contributes to the progressive MC truncation observed in dhomo; het mutants, we performed EdU incorporation assays in control and mutant larvae (**Fig. S2**). Consistent with interstitial growth, EdU labeling in controls frequently identified adjacent chondrocyte doublets within the same cellular layer, indicative of ongoing proliferation. In contrast, EdU-positive chondrocytes were rare in dhomo;het mutants and significantly less abundant compared than controls, suggesting reduced proliferative activity. This decrease is consistent with a shift in cell differentiation status that limits chondrocyte expansion and MC growth, particularly away from the proximal MC region where interstitial growth occurs and where reductions in cell number are evident by 14 dpf (**Fig. 4F**).

Our findings differ from previous studies reporting that Noggin deletion leads to mandibular thickening with apparent normal length, attributed to an increased number of chondrocytes within MC (Matsui and Klingensmith, 2014; Stottmann et al., 2001; Wang et al., 2013). To evaluate whether Noggin play a conserved role in chondrocytes, we analyzed noggin mutants in isolation. Wild-type, *nog2*^+/-^; *nog3*^+/-^, *nog2*^-/-^; *nog3*^+/-^ and *nog2*^-/-^; *nog3*^-/-^ larvae were stained with DAPI and chondrocytes were quantified (**Fig. S3A, B**). Compared with wildtype controls, all other genotypes exhibited a significant increase in chondrocyte number, consistent with a conserved role for BMP antagonists in restraining chondrocyte numbers and highlighting a robust dosage response of chondrocytes to BMP signaling. Although not statistically significant, we also noted an increasing trend in chondrocyte number as BMP antagonist alleles were progressively lost. Because *nog2*^-/-^; *nog3*^-/-^ mutants do not survive to adulthood, we next assessed the long-term consequences of reduced Noggin expression by examining adult *nog2*^-/-^; *nog3*^+/-^ fish. These mutants displayed a variable and incompletely penetrant phenotype, including mandibular truncation and splayed lower jaws (**Fig. S3C**). Quantification of lower jaw position relative to the upper jaw confirmed a reduction in mandibular length. Notably, this phenotype was less penetrant and less severe than that observed in *grem1a*^-/-^; *nog2*^-/-^; *nog3*^+/-^ mutants (**Fig. S3D**). Together, these findings support a conserved role for BMP antagonists in controlling chondrocyte number and reveal functional interactions between Gremlin and Noggin family antagonists in sustaining mandibular growth.

### Altered chondrocyte differentiation precedes MC truncation

Previous studies reported altered chondrocyte differentiation within the intermediate domain of Meckel’s cartilage in Noggin mutants (Wang et al., 2013). To determine whether disrupted differentiation contributes to the enlarged chondrocyte size observed in our mutants, a characteristic associated with hypertrophic chondrocytes, we examined markers of chondrocyte maturation by combinatorial RNAscope in situ hybridization (**Fig. 5A, B**). At 6 dpf in control zebrafish, the hyaline chondrocyte marker *col2a1a* was broadly expressed throughout MC, with reduced signal toward the distal domain (**Fig. 5A**). In contrast, the hypertrophic marker *col10a1a* was largely excluded from MC chondrocytes and instead localized to surrounding dermal bone. However, distal chondrocytes expressed the pre-hypertrophic marker *ihha* (**Fig. 5A**). In both dhomo; het and triple homo mutants, *col2a1a* expression was present within the proximal MC but reduced across more distal regions. Notably, *col10a1a* transcripts were ectopically detected within distal MC chondrocytes (**Fig. 5A**). Mutants also exhibited an expanded *ihha* expression domain that extended beyond regions marked by *col10a1a* and overlapped with *col2a1a*-positive territories (**Fig. 5A**). By 14 dpf, control MCs maintained *col2a1a* expression along most of the cartilage, with *col10a1a*⁺ chondrocytes restricted to the distal tip (**Fig. 5B**). In dhomo; het mutants, however, *col10a1a* expression domains were expanded and the reciprocal *col2a1a* domains were reduced (**Fig. 5B**). Consistent with these findings, *col2a1a*:mCherry reporter activity was diminished in mutants at 14 dpf, indicating impaired maintenance of the hyaline chondrocyte program (**Fig. S4**). Quantification of MC regions defined by *col2a1a*⁺, *col10a1a*⁺, or *ihha*^+^ chondrocytes confirmed a significant expansion of hypertrophic-like (*col10a1a*^+^ or *ihha*^+^) domains and a reduction in hyaline (*col2a1a*⁺) domains at both 6 dpf and 14 dpf (**Fig. 5C**). At 6 dpf, triple homozygous fish displayed a non-significant trend toward expanded hypertrophic markers and reduced hyaline cartilage markers, indicative of a dosage-sensitive response to BMP antagonist loss (**Fig. 5C**). Consistent with these observations, pSMAD1/5 activity showed a non-significant increase in the dhomo; het MC and a significant increase in triple homozygous zebrafish compared with controls, reinforcing the requirement for tight regulation of BMP signaling during chondrocyte maturation. (**Fig. S5**).

**Figure 5.**
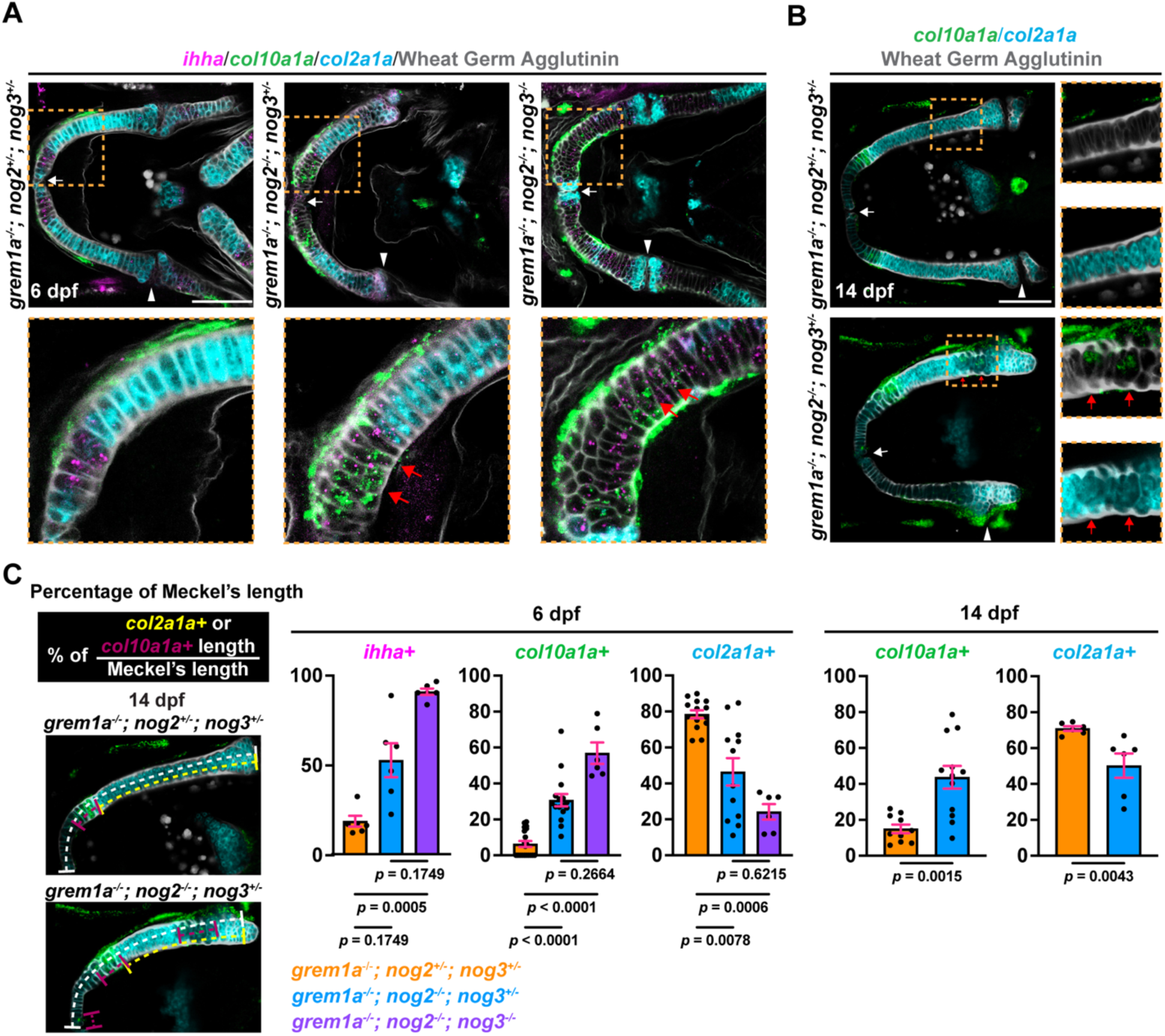
Loss of BMP antagonists disrupts normal chondrocyte differentiation along Meckel’s cartilage. (**A**) Single Z-plane confocal images of Meckel’s cartilage following in situ hybridization for *col2a1a* (blue), *col10a1a* (green), and *ihha* (magenta) counter-stained with Wheat Germ Agglutinin (gray), in control and mutant skeletons at 6 dpf. Yellow dashed boxes indicate regions shown in higher-magnification insets. (**B**) Single Z-plane confocal images of Meckel’s cartilage following in situ hybridization for *col2a1a* (blue) and *col10a1a* (green), counterstained with Wheat Germ Agglutinin (gray), in control and dhomo; het mutant skeletons at 14 dpf. Yellow dashed boxes indicate regions shown in higher-magnification insets. (**C**) Representative 14 dpf schematic of the quantification strategy and corresponding analysis of the percentage of *col2a1a* or *col10a1a* domains within control and mutant Meckel’s cartilage (n = 14 control, 12 dhomo; het, and 6 triple homo Meckel’s cartilages for *col2a1a*, n = 20 control, 16 dhomo; het, and 6 triple homo Meckel’s cartilages for *col10a1a* and n = 6 Meckel’s cartilages per genotype for *ihha* at 6 dpf; n = 6 control and dhomo; het Meckel’s cartilages for *col2a1a* and n = 10 control and 12 dhomo; het Meckel’s cartilages for *col10a1a* at 14 dpf). Quantification reveals expanded pre-hypertrophic and hypertrophic domains and a smaller *col2a1a+* domain in both dhomo; het and triple homo larval fish. White arrows point to the anterior of the Meckel’s cartilage and arrowheads indicated the posterior of the Meckel’s cartilage. Red arrows mark ectopic *col10a1a*+ chondrocytes. Scale bars, 100 µm. Error bars represent SEM.

To determine whether activation of BMP signaling in chondrocytes is sufficient to alter chondrocyte differentiation, we generated a *col2a1a*-R2-E1B:*CAbmpr1ba*-p2a-mCherry construct and injected it into wild-type embryos (Dale and Topczewski, 2011; Peskin et al., 2023). We then compared gene expression and cellular morphology between mCherry⁺ (harboring transgene) and mCherry⁻ (internal controls) chondrocytes. Expression of the construct was sufficient to induce ectopic activation of *ihha* and *col10a1a* (**Fig. 6A**). Quantification of chondrocyte volume revealed a significant increase in cell size among mCherry⁺ chondrocytes, consistent with acquisition of a hypertrophic-like state (**Fig. 6B**). In addition to these molecular and morphometric changes, mCherry⁺ chondrocytes exhibited instances of reduced organization and disrupted stacking, suggesting that elevated BMP signaling perturbs normal chondrocyte differentiation and tissue architecture within MC (**Fig. 6A**).

**Figure 6.**
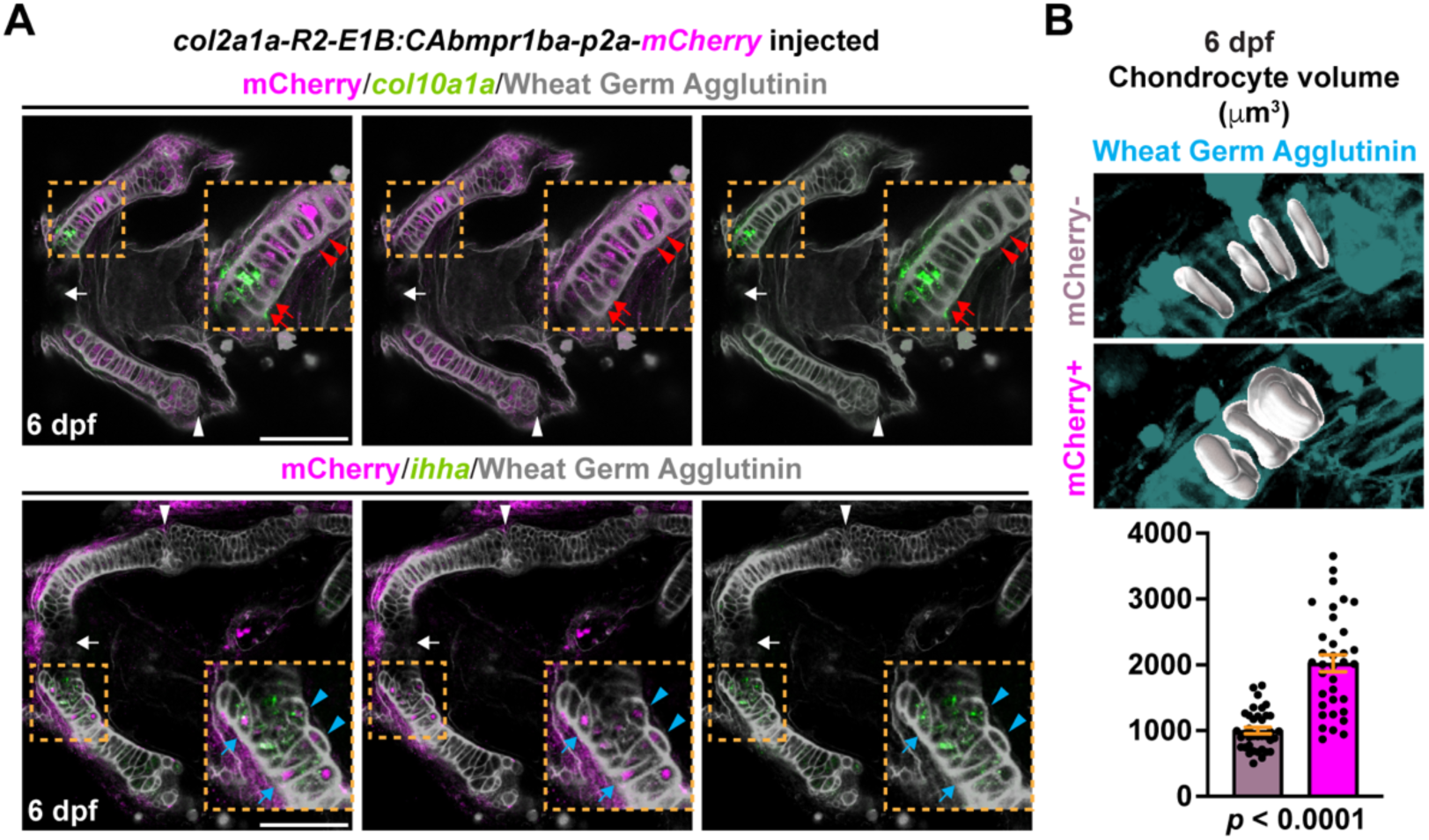
Activation of BMP signaling within chondrocytes is sufficient to induce a hypertrophic program and disrupt chondrocyte stacking. (**A**) Single Z-plane confocal images of Meckel’s cartilage following in situ hybridization for *col10a1a* (n = 7 zebrafish) or *ihha* (n = 4 zebrafish) (green), combined with immunostaining for mCherry (magenta) and staining for Wheat Germ Agglutinin (gray) in embryos injected with a *col2a1a-R2:CAbmpr1ba-p2a-mCherry* construct. Yellow dashed boxes indicate regions shown in higher-magnification insets. mCherry-positive cells, expressing constitutively active *bmpr1ba*, display ectopic expression of pre-hypertrophic (blue arrows) and hypertrophic markers (red arrows) and exhibit enlarged and disorganized chondrocyte organization (red arrowheads and blue arrowheads). (**B**) Representative 3D reconstructions of mCherry-positive and mCherry-negative chondrocytes in injected embryos. Quantification of chondrocyte volume (n = 35 positive or n = 37 negative chondrocytes from 14 animals) demonstrates increased cell size in mCherry-positive cells at 6 dpf. White arrows point to the anterior of the Meckel’s cartilage and arrowheads indicated the posterior of the Meckel’s cartilage. Scale bars, 100 µm. Error bars represent SEM.

### BMP antagonist loss disrupts regional skeletal mesenchyme but not early osteogenesis

Our in situ analyses suggest that BMP antagonists may also function within skeletal mesenchyme in addition to chondrocytes. We therefore next examined other skeletal cell populations associated with MC. Analysis of previously published single-cell RNA-sequencing datasets identified a *tnn*⁺ *postnb*⁺ population present by 5 dpf, with *tnn* marking a subset of the *postnb*⁺ mesenchyme that overlapped with BMP antagonist expression (**Fig. S1A**). Using combinatorial RNAscope in situ hybridization, we confirmed co-expression of *tnn* and *postnb* at 14 dpf. Double-positive cells were localized predominantly along the lateral aspect of MC, whereas *postnb*-single-positive cells were enriched proximally (**Fig. S6A**). Additional in situ analyses revealed co-expression of BMP antagonists within a subset of laterally located *tnn*⁺ mesenchymal cells (**Fig. S6B**). Proximal *tnn*⁺ mesenchyme was sandwiched in between MC and *col10a1a*⁺ dermal bones, while distal *tnn*⁺ mesenchyme expression was associated with low *col10a1a* expression (**Fig. S6C**). Interestingly, the *tnn*⁺ mesenchyme tapered off before reaching the *col10a1a*⁺ chondrocytes. We next evaluated the distribution of *tnn*⁺ mesenchymal cells and *ifitm5*⁺ osteoblasts at 14 dpf in control and mutant zebrafish. At this stage, MC is surrounded by three intramembranous bones: the dentary, anguloarticular, and retroarticular. All three bones were present in both control and mutant fish with normal morphology, although mutant dentary bone was shorter than controls (**Fig. S6D**). In contrast, although *tnn*⁺ mesenchyme remained normally distributed along the proximal and medial MC and within *tnn*⁺ tendons (Roberts et al., 2026), the distal *tnn*⁺ domain was selectively lost in mutants (**Fig. S6D**). This disruption is consistent with a requirement for BMP antagonists in establishing or maintaining region-specific skeletal mesenchymal identities.

Collectively, these data support a model in which precise modulation of BMP signaling regulates chondrocyte differentiation and skeletal mesenchyme development, thereby preventing ectopic hypertrophic-like differentiation. Disruption of BMP antagonism elevates pathway activity, driving aberrant chondrocyte differentiation, defective cellular organization, and impaired MC elongation. This failure of cartilage growth may be sufficient to drive secondary mandibular truncation as surrounding bones form around a reduced MC scaffold (**Fig. 7**).

**Figure 7.**
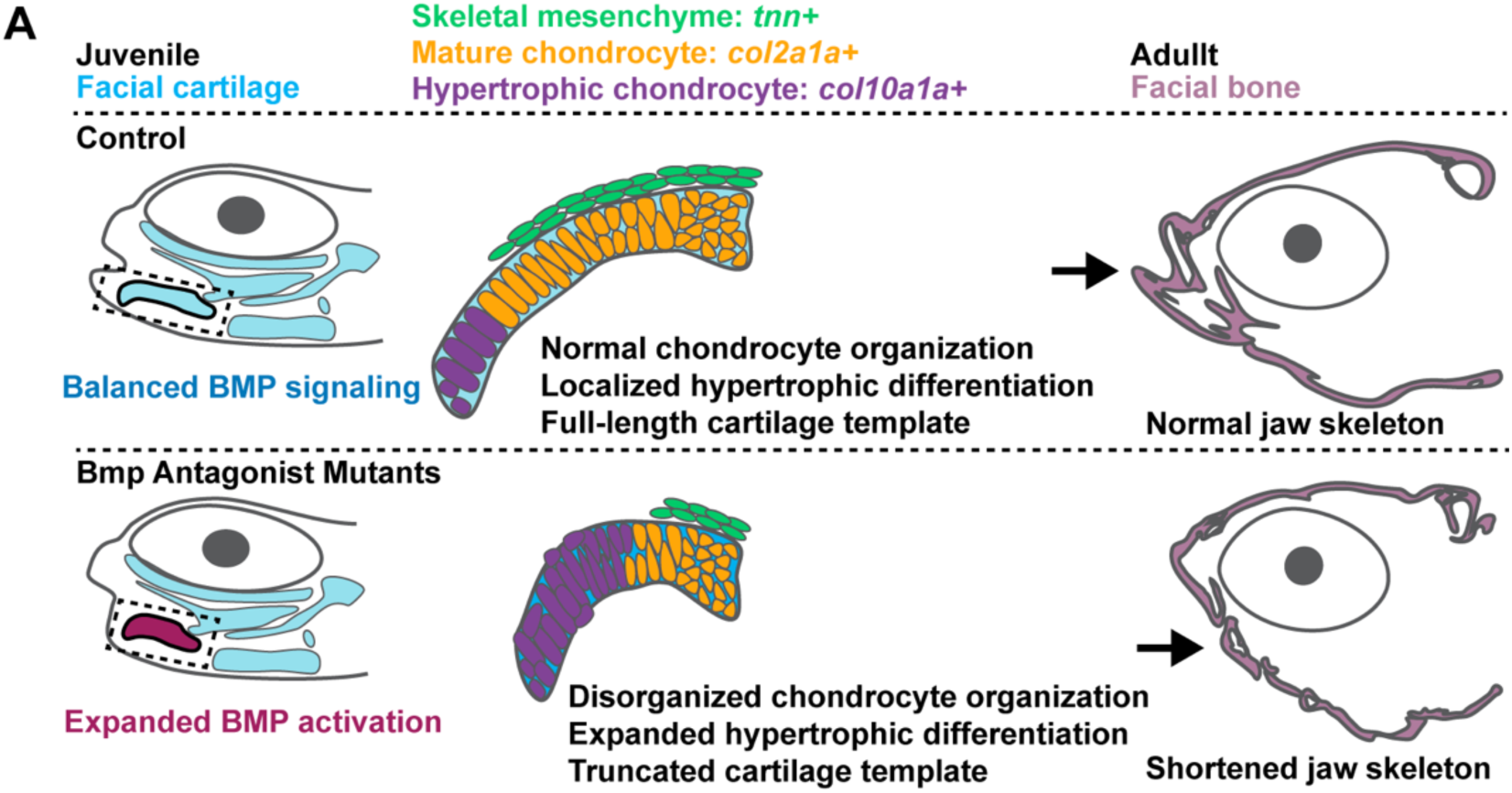
Model for sustained requirement for BMP antagonism. Summary model illustrating the cellular and molecular defects in compound mutant zebrafish and their contribution to mandible truncation observed in adult zebrafish.

## Discussion

The Meckel’s cartilage (MC) is a key regulator of mandibular development across vertebrates. In mammals, the fate of MC is transient, with the cartilage ultimately lost shortly after birth. Mandibular growth then proceeds primarily through osteogenesis (Funato et al., 2009; Guo et al., 2023). Thus, in mammals, MC provides an initial template essential for establishing proper mandible size but is dispensable for patterning mandibular bones (Mori-Akiyama et al., 2003). While MC function has been extensively investigated in mammalian systems, the developmental and functional significance of a persistent MC in other vertebrates remains less well defined. In birds, fish, and many non-mammalian vertebrates, MC is retained and is thought to maintain its role as a structural scaffold for mandibular development, although this function has rarely been directly tested. In zebrafish, MC arise from the first pharyngeal arch and forms a central component of the cartilaginous lower jaw (Dougherty et al., 2012). Initially organized as bilateral cartilages that meet at the symphysis, MC becomes surrounded by dermal bones beginning as early as 4 dpf (Cubbage and Mabee, 1996; Eames et al., 2013). During early larval stages, MC growth has generally been attributed to chondrocyte intercalation, with minimal evidence of proliferation or changes in chondrocyte number between 4–6 dpf, despite increases in MC length (Chen et al., 2023; LeClair et al., 2009; Shwartz et al., 2012). However, proliferation may contribute again at later larval stages, as chondrocyte doublets within a single layer are observed by 14 dpf, consistent with interstitial growth (Eames et al., 2013). By adulthood, distal MC chondrocytes adjacent to the mandibular symphysis have likely undergone endochondral ossification, whereas intermediate and proximal regions persist, embedded within dermal bone (Wang et al., 2012). Notably, MC growth appears proportional to mandibular expansion, suggesting a sustained developmental coupling between cartilage and bone that extends beyond the transient template role observed in mammals.

Here, we investigated how BMP antagonists regulate MC development and mandibular growth. Prior studies demonstrated that Noggin restricts chondrocyte proliferation, and we find that loss of *nog2* and *nog3* similarly results in increased chondrocyte number, consistent with a conserved role for BMP antagonism in modulating chondrocyte cell number (Wang et al., 2013). In mice, this expanded cartilage template is associated with mandibular thickening through hypertrophic differentiation and endochondral ossification, reinforcing the framework that MC influences final mandibular morphology. However, because *nog2*^-/-^; *nog3*^-/-^ zebrafish mutants do not survive to adulthood, we were unable to assess long-term mandibular consequences in this genetic background. Instead, partial reduction of Noggin dosage (*nog2*^-/-^; *nog3*^+/-^) produced a variable and incompletely penetrant mandibular phenotype, including truncated or splayed jaws. These findings suggest that while BMP antagonists regulate chondrocyte proliferation across species, the morphological outcomes of antagonist loss diverge between mammals and vertebrates that retain MC.

To generate a more penetrant model, we leveraged prior evidence of Gremlin–Noggin interactions during zebrafish cranial suture formation, along with transcriptional datasets indicating overlapping antagonist expression during craniofacial development. Previous studies also proposed functional redundancy among BMP antagonists, particularly between Noggin and Chordin, where combined disruption leads to mandibular dysplasia (Stottmann et al., 2001). Consistent with cooperative antagonistic regulation, *grem1a*^-/-^; *nog2*^-/-^; *nog3*^+/-^ mutants exhibited fully penetrant mandibular truncations. Interestingly, at 6 dpf these compound mutants did not display increased chondrocyte number compared to *grem1a*^-/-^; *nog2*^+/-^; *nog3*^+/-^ controls, indicating that early changes in chondrocyte number cannot explain later mandibular phenotypes. In triple homozygous zebrafish, MC remained poorly extended with abnormal cartilage shape despite an increased chondrocyte number and cartilage volume, indicating that additional cellular defects contribute to reduced lower jaw size. Instead, our data identify disrupted chondrocyte differentiation as the most salient cellular defect, similar to those previously observed in mice (Wang et al., 2013). We show that hypertrophic differentiation programs normally arise within distal MC regions that neighbor the symphysis, detectable by 6 dpf and more pronounced by 14 dpf. These domains spatially coincide with regions where chondrocytes are replaced by bone to separate the mandibular symphysis and the persisting bilateral cartilage rods. In mutants, hypertrophic markers were expanded and ectopically activated, consistent with altered maturation. Differentiation defects were most apparent away from the proximal end of MC, where *col2a1a* was retained in both dhomo; het and triple homo zebrafish. By 14 dpf, local quantification in this region revealed selective reductions in cell number, coinciding with the domain where interstitial growth emerges at later larval stages and where proliferative chondrocytes are reduced in mutants. Although transcriptional diversity within zebrafish MC remains incompletely characterized, these findings imply the existence of region-specific regulatory programs. Notably, truncations were already apparent in triple homozygous mutants by 6 dpf, suggesting that reduced BMP antagonist dosage accelerates cellular and molecular defects that lead to early lower jaw truncation.

BMP antagonists were also expressed within a subset of skeletal mesenchyme that exhibited sensitivity to the loss of BMP antagonism. This mesenchymal population resides adjacent to hyaline cartilage but is spatially separated from the hypertrophic zone, hinting at a possible function to support local suppression of BMP signaling and thereby prevent ectopic chondrocyte differentiation. Ultimately, mutants exhibited widespread cellular features consistent with hypertrophic-like differentiation, including enlarged chondrocyte volume. These cellular alterations were accompanied by pronounced disruption of cartilage architecture. By 14 dpf, mutant MCs lacked the characteristic rod-like morphology, instead appearing irregular and poorly organized. Similar defects were induced by constitutive activation of *Bmpr1a* in chondrocytes, linking elevated BMP signaling directly to aberrant differentiation and tissue disorganization. These observations parallel findings in chick embryos, where caBMPR activation disrupted chondrocyte stacking and cartilage organization, supporting a conserved mechanistic relationship between BMP signaling and cartilage differentiation dynamics (Ashique et al., 2002).

A striking conclusion from this work is the divergence between mammalian and zebrafish phenotypic outcomes following BMP pathway activation. In mice, enhanced BMP signaling results in mandibular thickening. In zebrafish, early skeletal patterning remains largely intact (particularly in *grem1a*^-/-^; *nog2*^-/-^; *nog3*^+/-^ fish), yet later MC growth is selectively impaired, leading to a shortened MC and likely, as a consequence, a truncated mandibular. This distinction likely reflects fundamental differences in MC function across species. In mammals, the transient nature of MC limits its long-term contribution to mandibular growth. In contrast, zebrafish MC persists and must remain developmentally coordinated with surrounding bone. Under these conditions, sustained disruption of chondrocyte differentiation compromises cartilage elongation, organization, and proliferative capacity, ultimately restricting mandibular growth. Our findings therefore support a model in which MC functions not merely as an early template in zebrafish but as a persistent scaffold required for continued mandibular expansion. BMP antagonists play a critical role in preserving this function by restraining ectopic hypertrophic differentiation and maintaining cartilage architecture. When BMP antagonism is reduced, elevated BMP signaling drives aberrant chondrocyte maturation, disrupts tissue organization, and limits sustained cartilage growth. Because intramembranous bones form around a reduced MC framework, mandibular size cannot be restored. Indeed, osteogenesis exhibited normal timing and organization in mutant zebrafish at 14 dpf. However, because these mutations are global knockout alleles, we cannot exclude the possibility that later disruptions in mandibular osteogenesis further contribute to the mandibular phenotype. In addition, BMP antagonists may have a transient role in the MC that produces lasting consequences, rather than functioning continuously throughout mandibular development. Indeed, in mice, although Noggin remains expressed in the MC, BMP ligands are sharply downregulated (Wang et al., 2013). However, parallels with avian models further reinforce this interpretation of BMP signaling requirements in the lower jaw. Constitutive BMPR activation in chick embryos similarly produces mandibular truncation phenotypes, suggesting that persistent MC-bearing vertebrates share a heightened dependence on balanced BMP signaling to sustain cartilage-driven jaw growth (Ashique et al., 2002). Future studies that enable spatiotemporal control of BMP antagonist deletion will further refine our understanding of these processes.

Collectively, this study reveals conserved cellular requirements of BMP antagonism with species-specific consequences and highlights the prolonged functional requirement of Meckel’s cartilage in non-mammalian mandibular development. These findings underscore the importance of antagonistic BMP regulation in coordinating cartilage differentiation, tissue organization, and skeletal growth.

## Methods

### Zebrafish husbandry

All experiments were approved by the Institutional Animal Care and Use Committee at the University of California, Los Angeles (Protocol #ARC-2022-044). Published lines included *Tg(col2a1aBAC:mCherry-NTR)^el559^* (Askary, Dev Cell, 2015), *grem1a ^el1049^, nog2 ^el1050^*, and *nog3 ^el1051^* (Farmer, Nature Comm, 2024).

### Constitutively active form of bmpr1ba (CAbmpr1ba) cloning and injection

Constitutively active (CA) form of *bmpr1ba*, *CAbmpr1ba-p2a-mCherry (Peskin et al., 2023)*, driven by *col2a1a-R2* enhancer-E1B minimum promoter (Dale and Topczewski, 2011) was inserted into the construct flanked with Tol2 sequence through In-Fusion Snap Assembly (Cat no. 638946). One-cell stage embryos were co-injected with *col2a1a-R2-E1B:CAbmpr1ba-p2a-mCherry* construct (20 ng/ul) with Tol2 transposase RNA (30 ng/ul).

### Skeletal preparations

Adult skeletal staining of bone and cartilage was performed as previously described (Farmer, Nature Comm, 2024). Embryonic (6 and 14 dpf) skeletal preparations were performed as previously described (Chen et al., 2023). All skeletal preparations were transferred through a glycerol gradient to 100% glycerol for imaging and dissection.

### Nuclei and cartilage ECM staining

For staining of chondrocyte nuclei or ECM, zebrafish heads were washed and permeabilized by 1% Triton-X in 1X PBS for several hours at room temperature. Samples with then incubated in 1% Triton-X/PBS with DAPI or Wheat Germ Agglutinin (10ug/ml, WGA-Alexa Fluor Plus 405, W56132; WGA-Alexa Fluor 488, W11261; WGA-Alexa Fluor 647, W32466, Thermo Fisher Scientific) at least overnight at 4°C in dark.

### mRNA in situ hybridization

Zebrafish embryos were fixed with 4% PFA/1X PBS overnight at 4°C and heads were dissected for in situ hybridization. Detection of mRNA expression was performed through RNAscope Multiplex Fluorescent Reagent Kit v2 Assay (Advanced Cell Diagnostics) per manufacturer’s instructions on whole zebrafish embryos with modifications. After rehydration through 1X PBST (0.1% Tween-20)/methanol series, zebrafish heads were treated with 3% hydrogen peroxide for 10 mins at room temperature. After washing with PBST (added 1% BSA), heads were treated with Target Retrieval for 15 mins at 40°C. Permeabilization was achieved using Protease Plus for 10 mins (6 dpf) or 15 mins (14 dpf) at 40°C. The following probes were used: Channel 1, dr-*grem1a*, dr-*ihha*, dr-*postnb*, dr-*ifitm5*; Channel 2, dr-*col2a1a*, dr-*tnn*; Channel 3, dr-*nog3*, dr-*col10a1a*; Channel 4, dr-*nog2*. After signal amplification steps (AMP1-3), zebrafish heads were pretreated with HRP blocker for 15 mins at 40°C preceding to HRP steps. For counter-staining of nuclei or cartilage ECM, DAPI or Wheat Germ Agglutinin was added into Wash Buffer and incubated for at least overnight at 4°C in dark.

### Immunofluorescence

For detection of mCherry after RNAscope in situ hybridization, embryos were washed and blocked for 3 hr in 2% goat serum at room temperature, stained with primary antibody (1:200 anti-mCherry, Novus NBP2-25157) overnight and secondary antibody (1:300 Alexa Fluor 568 anti-rabbit, Thermo Fisher Scientific A-11011) overnight at 4°C. For phospho-Smad1/5 staining, embryos were fixed as described above and washed with TBS. Embryos were permeabilized with acetone for 30 mins in −20°C followed by RNAscope Protease Plus treatment for 30 mins at 40°C, and blocked for 3hr in 2% goat serum in TBDTx (1% BSA, 1% DMSO, 1% Triton-X in 1X TBS, pH 7.3) at room temperature. Heads were stained with primary antibody (1:200, anti-pSmad-1/5 (S463/465), Cell Signaling 9516) overnight and secondary antibody (1:500 Alexa Fluor 488 anti-rabbit, Thermo Fisher Scientific A-11008) overnight. DAPI or Wheat Germ Agglutinin was added along with secondary antibodies.

### EdU labeling

EdU (500 uM, Biosynth NE08701) was injected into the heart at 13 dpf and fish were incubated for 20 hours to allow for EdU incorporation. Zebrafish heads were then fixed with 4% PFA for 30 mins at room temperature, washed with 1% Triton-X/1X PBS, and permeabilized with RNAscope Protease Plus for 30 mins at 40°C followed by wash with 3% BSA in 1X PBS. To develop EdU signal, heads were incubated with 8 uM Sulfo-Cyanine 3 Azide (Lumiprobe D1330) /100 mM Tris-HCl 7.6/4 mM CuSo4/100 mM sodium ascorbate (made fresh) for 1 hr at room temperature. Heads were washed with 3% BSA in 1X PBS followed by a wash with 1% Triton-X/1X PBS, and then incubated with DAPI in 1% Triton-X/1X PBS overnight at 4°C in dark for counter-staining of nuclei.

### Imaging

Brightfield images and movies of adult zebrafish and skeletal preparations were imaged and dissected under the LEICA S9i stereo microscope. All confocal images were taken by Zeiss LSM980 confocal microscope with ZEN software.

### microCT

Representative samples (both mutant and size-matched control) were scanned using a Quantum GX2 micro-CT Imaging System (Revvity, Hopkinton, MA, USA). Whole-body scans were conducted with the following settings: isotropic voxel size of 10 μm, 36 mm field of view, voltage of 50 kV, current of 155 μA, and no filter. Subsequent skull-only imaging was conducted with the following settings: isotropic voxel size of 5μm, 5 mm field of view, voltage of 50 kV, current of 155 μA, and no filter. Three-dimensional reconstructions were visualized with AnalyzeDirect 14.0.

### Quantification

Lower jaw extension was assessed by measuring the horizontal distance from the anterior tip of the lower jaw to the anterior tip of neurocranium (6 and 14 dpf) or to the anterior tip of the upper jaw (adult) in ImageJ. DAPI+ chondrocytes within Meckel’s cartilage were manually counted through confocal Z-stacks in ImageJ for 6 dpf embryos, and counted by the Spot detection function in Imaris for 14 dpf juveniles. Chondrocytes were identified by the morphology and orientation of the nuclei and outlined by Wheat Germ Agglutinin staining. The number of EdU+/DAPI+ chondrocytes in Meckel’s cartilage were manually counted across confocal z stacks in ImageJ. Meckel’s cartilage was outlined by Wheat Germ Agglutinin staining and traced across Z-stacks in Imaris that allowed 3D reconstruction by the Surface detection function and 3D rendering to measure the volume of Meckel’s cartilage. The cross-section perpendicular to the Meckel’s cartilage at the middle point of the cartilage was used to measure dorsal-ventral and lateral-medial diameters of the Meckel’s cartilage in ImageJ. To calculate the cell volume of chondrocytes using Imaris in controls and mutants, the largest chondrocyte at the middle point of the Meckel’s cartilage and two neighboring cells at the both sides were selected. Cell outlines stained by Wheat Germ Agglutinin were traced across the Z-stacks to construct the 3D surface of chondrocytes and used to measure chondrocyte volumes. To measure chondrocyte volumes in embryos injected with the constitutively active form of *bmpr1ba*, cells with clear mCherry signal were selected and cells negative for mCherry were used as controls. Lengths of expression along Meckel’s cartilage was assessed by drawing a line along the middle of Meckel’s cartilage on 3D sections across the Meckel’s cartilage compared to the total length to calculate the percentage of Meckel’s length covered by selected genes and phospho-Smad1/5 using ImageJ. Statistical analyses were calculated by ANOVA followed with Kruskal-Wallis test for comparison between three and more groups and by Mann-Whitney test for comparison between two groups in Prism. Error bars represent the SEM.

## Contributions

H.C. and D.T.F conceived and designed the study. J.D. performed experiments. P.X. provided initial characterization of noggin phenotypes. T.L. performed microCT experiments. H.C performed experiments and bioinformatic analysis. D.T.F. supervised the research. H.C. and D.T.F. wrote the manuscript.

## Acknowledgements

The authors thank the Sagasti, Chen and Tornini labs for the feedback during shared lab meetings. The authors also thank Michel Bagnat for sharing the *CAbmpr1ba-p2a-mCherry* construct.

## Funding

HHMI Hanna H. Gray Fellows Program (D.T.F), HHMI Freeman Hrabowski Program (D.T.F.)

## Competing interests

The authors have no competing interests to declare.

---

**Movie 1.** Adult mutant fish have a truncated lower jaw with an intact jaw joint (n = 4 fish per genotype). Control shown from 0-6 seconds and dhomo: hets shown from 7-12 seconds.

## Notes

### Competing Interest Statement

The authors have declared no competing interest.

## References

1. Adel Al-Lami, H., Barrell, W.B., Liu, K.J., 2016. Micrognathia in mouse models of ciliopathies. Biochem Soc Trans 44, 1753–1759.

2. Anthwal, N., Joshi, L., Tucker, A.S., 2013. Evolution of the mammalian middle ear and jaw: adaptations and novel structures. J Anat 222, 147–160.

3. Ashique, A.M., Fu, K., Richman, J.M., 2002. Signalling via type IA and type IB bone morphogenetic protein receptors (BMPR) regulates intramembranous bone formation, chondrogenesis and feather formation in the chicken embryo. Int J Dev Biol 46, 243–253.

4. Bonatto Paese, C.L., Brooks, E.C., Aarnio-Peterson, M., Brugmann, S.A., 2021. Ciliopathic micrognathia is caused by aberrant skeletal differentiation and remodeling. Development 148.

5. Bonilla-Claudio, M., Wang, J., Bai, Y., Klysik, E., Selever, J., Martin, J.F., 2012. Bmp signaling regulates a dose-dependent transcriptional program to control facial skeletal development. Development 139, 709–719.

6. Chai, Y., Jiang, X., Ito, Y., Bringas, P., Jr., Han, J., Rowitch, D.H., Soriano, P., McMahon, A.P., Sucov, H.M., 2000. Fate of the mammalian cranial neural crest during tooth and mandibular morphogenesis. Development 127, 1671–1679.

7. Chen, H.J., Barske, L., Talbot, J.C., Dinwoodie, O.M., Roberts, R.R., Farmer, D.T., Jimenez, C., Merrill, A.E., Tucker, A.S., Crump, J.G., 2023. Nuclear receptor Nr5a2 promotes diverse connective tissue fates in the jaw. Dev Cell 58, 461–473 e467.

8. Chen, Y., Wang, Z., Chen, Y., Zhang, Y., 2019. Conditional deletion of Bmp2 in cranial neural crest cells recapitulates Pierre Robin sequence in mice. Cell Tissue Res 376, 199–210.

9. Cubbage, C.C., Mabee, P.M., 1996. Development of the cranium and paired fins in the zebrafish Danio rerio (Ostariophysi, Cyprinidae). J Morphol 229, 121–160.

10. Dale, R.M., Topczewski, J., 2011. Identification of an evolutionarily conserved regulatory element of the zebrafish col2a1a gene. Dev Biol 357, 518–531.

11. DeLaurier, A., 2019. Evolution and development of the fish jaw skeleton. Wiley Interdiscip Rev Dev Biol 8, e337.

12. Dougherty, M., Kamel, G., Shubinets, V., Hickey, G., Grimaldi, M., Liao, E.C., 2012. Embryonic fate map of first pharyngeal arch structures in the sox10: kaede zebrafish transgenic model. J Craniofac Surg 23, 1333–1337.

13. Eames, B.F., DeLaurier, A., Ullmann, B., Huycke, T.R., Nichols, J.T., Dowd, J., McFadden, M., Sasaki, M.M., Kimmel, C.B., 2013. FishFace: interactive atlas of zebrafish craniofacial development at cellular resolution. BMC Dev Biol 13, 23.

14. Fabian, P., Tseng, K.C., Thiruppathy, M., Arata, C., Chen, H.J., Smeeton, J., Nelson, N., Crump, J.G., 2022. Lifelong single-cell profiling of cranial neural crest diversification in zebrafish. Nat Commun 13, 13.

15. Farmer, D.T., Dukov, J.E., Chen, H.J., Arata, C., Hernandez-Trejo, J., Xu, P., Teng, C.S., Maxson, R.E., Crump, J.G., 2024. Cellular transitions during cranial suture establishment in zebrafish. Nat Commun 15, 6948.

16. Funato, N., Chapman, S.L., McKee, M.D., Funato, H., Morris, J.A., Shelton, J.M., Richardson, J.A., Yanagisawa, H., 2009. Hand2 controls osteoblast differentiation in the branchial arch by inhibiting DNA binding of Runx2. Development 136, 615–625.

17. Guo, J., Yu, S., Zhang, H., Zhang, L., Yuan, G., Liu, H., Chen, Z., 2023. Klf4 haploinsufficiency in Sp7+ lineage leads to underdeveloped mandibles and insufficient elongation of mandibular incisor. Biochim Biophys Acta Mol Basis Dis 1869, 166636.

18. Hu, D., Colnot, C., Marcucio, R.S., 2008. Effect of bone morphogenetic protein signaling on development of the jaw skeleton. Dev Dyn 237, 3727–3737.

19. Iwaya, C., Suzuki, A., Iwata, J., 2023. Loss of Sc5d results in micrognathia due to a failure in osteoblast differentiation. J Adv Res.

20. LeClair, E.E., Mui, S.R., Huang, A., Topczewska, J.M., Topczewski, J., 2009. Craniofacial skeletal defects of adult zebrafish Glypican 4 (knypek) mutants. Dev Dyn 238, 2550–2563.

21. Liu, W., Selever, J., Murali, D., Sun, X., Brugger, S.M., Ma, L., Schwartz, R.J., Maxson, R., Furuta, Y., Martin, J.F., 2005. Threshold-specific requirements for Bmp4 in mandibular development. Dev Biol 283, 282–293.

22. Matsui, M., Klingensmith, J., 2014. Multiple tissue-specific requirements for the BMP antagonist Noggin in development of the mammalian craniofacial skeleton. Dev Biol 392, 168–181.

23. Merrill, A.E., Eames, B.F., Weston, S.J., Heath, T., Schneider, R.A., 2008. Mesenchyme-dependent BMP signaling directs the timing of mandibular osteogenesis. Development 135, 1223–1234.

24. Mori-Akiyama, Y., Akiyama, H., Rowitch, D.H., de Crombrugghe, B., 2003. Sox9 is required for determination of the chondrogenic cell lineage in the cranial neural crest. Proc Natl Acad Sci U S A 100, 9360–9365.

25. Parada, C., Han, D., Grimaldi, A., Sarrion, P., Park, S.S., Pelikan, R., Sanchez-Lara, P.A., Chai, Y., 2015. Disruption of the ERK/MAPK pathway in neural crest cells as a potential cause of Pierre Robin sequence. Development 142, 3734–3745.

26. Peskin, B., Norman, J., Bagwell, J., Lin, A., Adhyapok, P., Di Talia, S., Bagnat, M., 2023. Dynamic BMP signaling mediates notochord segmentation in zebrafish. Curr Biol 33, 2574–2581 e2573.

27. Roberts, R.R., Bhojwani, A., Tseng, K.C., Elliott, K., Chen, H.J., Teubner, L., Sherwood, D., Smeeton, J., Miller, C.L., Nayak, P.K., Subramanian, A., Schilling, T.F., Merrill, A.E., Crump, J.G., 2026. Gene regulatory programs underlying diversification of facial ligaments and tendons in zebrafish. Development 153.

28. Schlombs, K., Wagner, T., Scheel, J., 2003. Site-1 protease is required for cartilage development in zebrafish. Proc Natl Acad Sci U S A 100, 14024–14029.

29. Shwartz, Y., Farkas, Z., Stern, T., Aszodi, A., Zelzer, E., 2012. Muscle contraction controls skeletal morphogenesis through regulation of chondrocyte convergent extension. Dev Biol 370, 154–163.

30. Stottmann, R.W., Anderson, R.M., Klingensmith, J., 2001. The BMP antagonists Chordin and Noggin have essential but redundant roles in mouse mandibular outgrowth. Dev Biol 240, 457–473.

31. Svandova, E., Anthwal, N., Tucker, A.S., Matalova, E., 2020. Diverse Fate of an Enigmatic Structure: 200 Years of Meckel’s Cartilage. Front Cell Dev Biol 8, 821.

32. Wang, X., He, H., Tang, W., Zhang, X.A., Hua, X., Yan, J., 2012. Two origins of blastemal progenitors define blastemal regeneration of zebrafish lower jaw. PLoS One 7, e45380.

33. Wang, Y., Zheng, Y., Chen, D., Chen, Y., 2013. Enhanced BMP signaling prevents degeneration and leads to endochondral ossification of Meckel’s cartilage in mice. Dev Biol 381, 301–311.

34. Yoon, B.S., Ovchinnikov, D.A., Yoshii, I., Mishina, Y., Behringer, R.R., Lyons, K.M., 2005. Bmpr1a and Bmpr1b have overlapping functions and are essential for chondrogenesis in vivo. Proc Natl Acad Sci U S A 102, 5062–5067.

