## Supplemental Data for "BMP antagonism is required for mandible outgrowth in zebrafish"

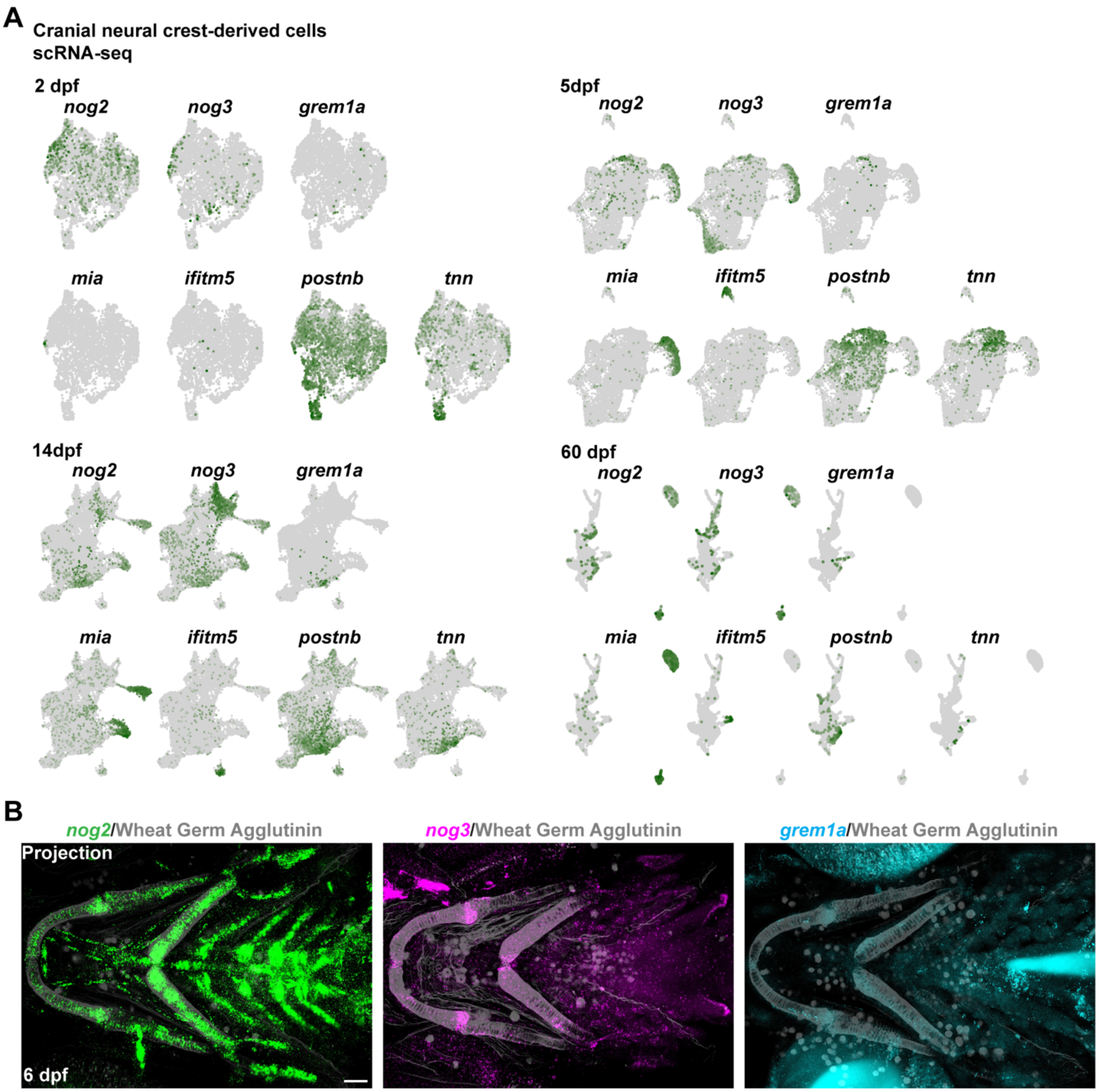


**Supp. Figure 1. BMP antagonists are expressed in chondrocytes and skeletal mesenchyme across craniofacial development.** (**A**) Analysis of BMP antagonists *nog2*, *nog3* and *grem1a* and skeletal markers *mia*, *ifitm5* and *tnn* using published single-cell RNA sequencing datasets from cranial neural crest-derived cells across development stages (Fabian et al., 2021). (**B**) Maximum projection confocal images of lower jaw skeleton following in situ hybridization for *nog2* (green), *nog3* (magenta) and *grem1a* (blue) counterstained with Wheat Germ Agglutinin (gray) at 6 dpf (n = 4 embryos for *nog2* and *nog3*; n = 2 embryos for *grem1a*) Scale bars, 50 μm.

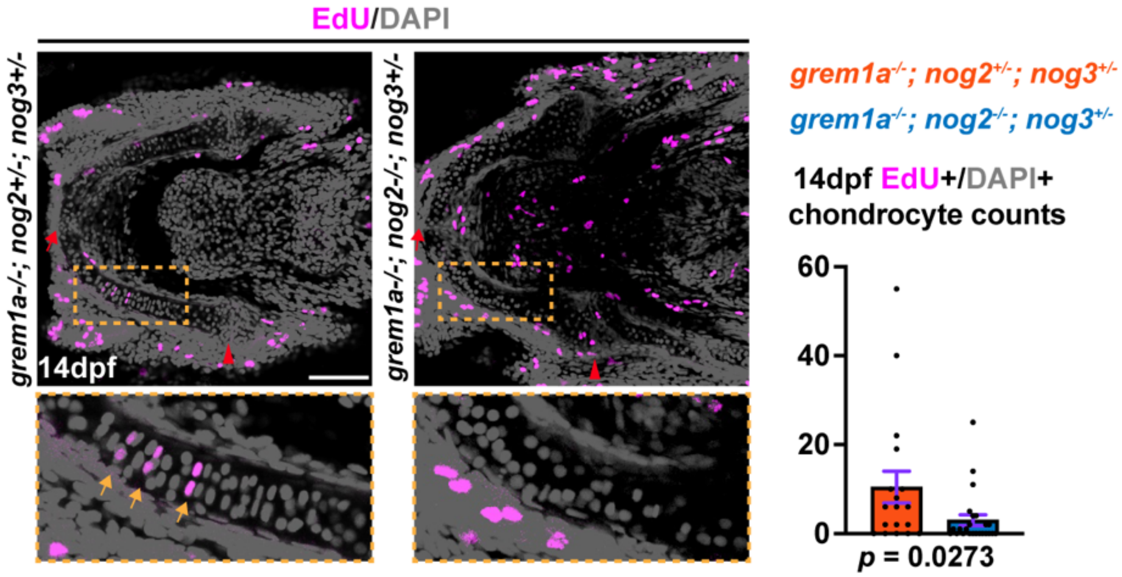


**Supp Figure 2. Chondrocytes are less proliferative in BMP antagonist mutants.** Representative single Z-plane confocal images of Meckel's cartilage following EdU incorporation from 13 to 14 dpf. Yellow dashed boxes indicate regions shown in higher-magnification insets. Yellow arrows denote EdU-positive chondrocytes. Red arrows point to the anterior of the Meckel's cartilage and arrowheads indicated the posterior of the Meckel's cartilage. Quantification of the number of EdU positive chondrocytes within a Meckel's cartilage showed reduced proliferation in dhomo; het mutants (n = 18 control and n = 24 dhomo; het Meckel's cartilages). Scale bars, 100 μm. Error bars represent SEM.

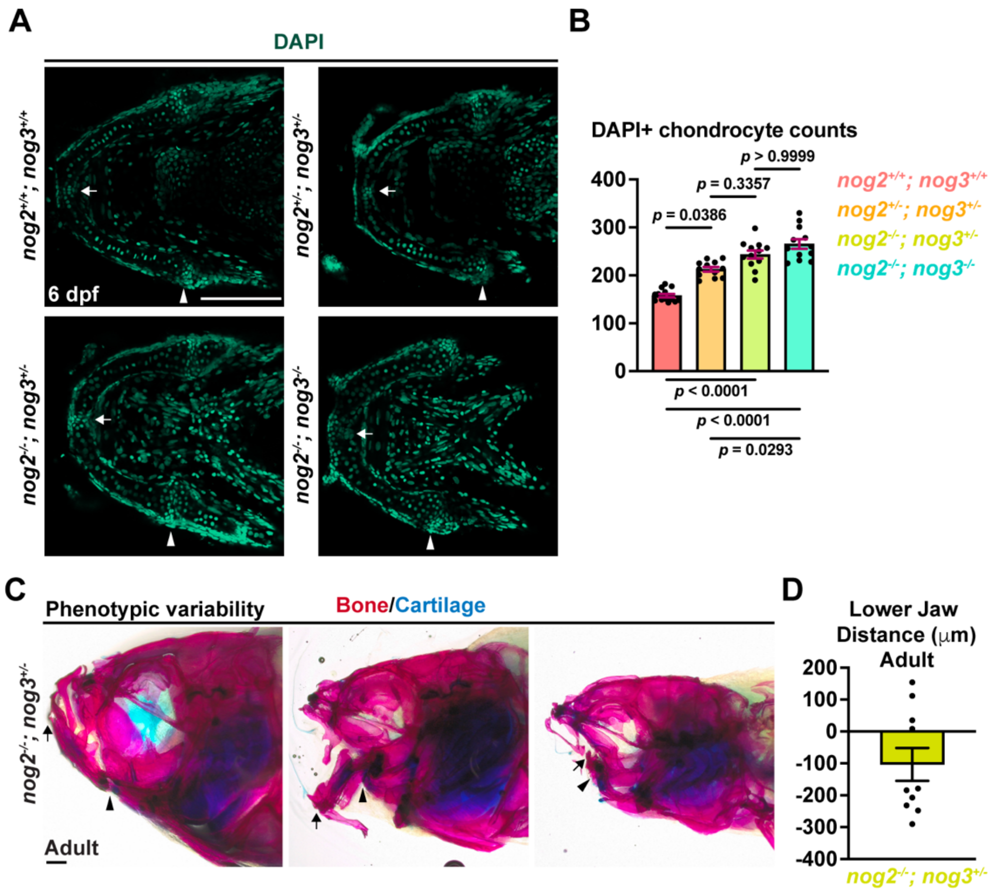


**Supp. Figure 3. Noggin genes exhibit a conserved requirement for chondrocyte number and display incompletely penetrant mandible truncations.** (**A**) Representative images of single Z-plane confocal images of DAPI stained wildtype, *nog2*$^{+/-}$; *nog3*$^{+/-}$, *nog2*$^{-/-}$; *nog3*$^{+/-}$ and *nog2*$^{-/-}$; *nog3*$^{-/-}$ zebrafish at 6 dpf. (**B**) Quantification of DAPI+ chondrocyte numbers across full confocal z-stacks of Meckel's cartilage indicates that total chondrocyte number is increased in all conditions compared to wildtype (n = 14 wildtype and n = 12 for other genotypes Meckel's cartilages). **(C)** Skeletal staining of bone (red) and cartilage (blue) of adult *nog2*$^{-/-}$; *nog3*$^{+/-}$ mutants (n =10) reveal partial mandible truncations. Arrows point to the anterior of the Meckel's cartilage and arrowheads indicated the posterior of the Meckel's cartilage. Scale bars, (A) 100 μm, (C) 500 μm. Error bars represent SEM.

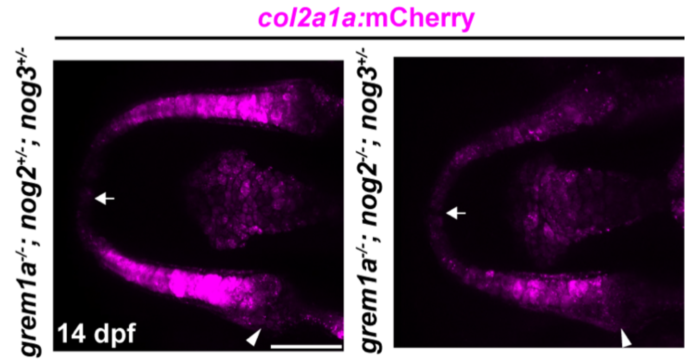


**Fig. S4. Reduced col2a1a enhancer activity in BMP antagonist mutants.** Maximum projection confocal images of control and dhomo; het Meckel's cartilage carrying a *col2a1a:mCherryNTR* reporter (n = 5 per genotype). mCherry signal is diminished in dhomo; het mutant cartilages. White arrows point to the anterior of the Meckel's cartilage and arrowheads indicated the posterior of the Meckel's cartilage. Scale bars, 100 µm.

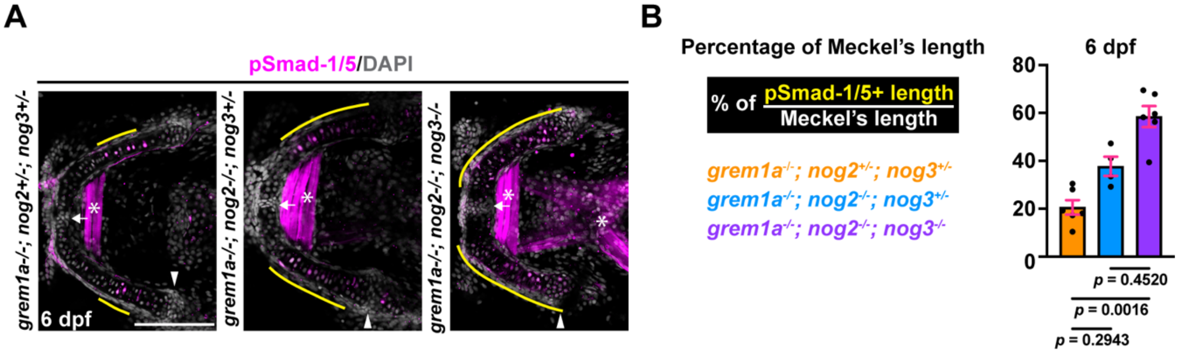


**Supp. Figure 5. pSmad+ chondrocytes are expanded in mutant Meckel's cartilages.** Single Z-plane confocal images of Meckel's cartilage immunostained for pSmad1/5 (magenta) counter-stained with DAPI (gray) reveal an expanded domain of pSmad+ chondrocytes (outlined by the outer yellow line) in *grem1a*$^{-/-}$; *nog2*$^{-/-}$; *nog3*$^{+/-}$ , with further expansion in *grem1a*$^{-/-}$; *nog2*$^{-/-}$; *nog3*$^{-/-}$ zebrafish at 6 dpf (n = 6 control, 4 dhomo; het and 6 triple homo Meckel's cartilages). (**B**) Quantification of the fraction of pSmad1/5 signal within MC reveals a non-significant expansion in *grem1a*$^{-/-}$; *nog2*$^{-/-}$; *nog3*$^{+/-}$ fish and a significant increase in *grem1a*$^{-/-}$; *nog2*$^{-/-}$; *nog3*$^{-/-}$ fish compared to controls. White arrows point to the anterior of the Meckel's cartilage and arrowheads indicated the posterior of the Meckel's cartilage. Asterisks mark background single in muscle. Scale bars, 100 µm.

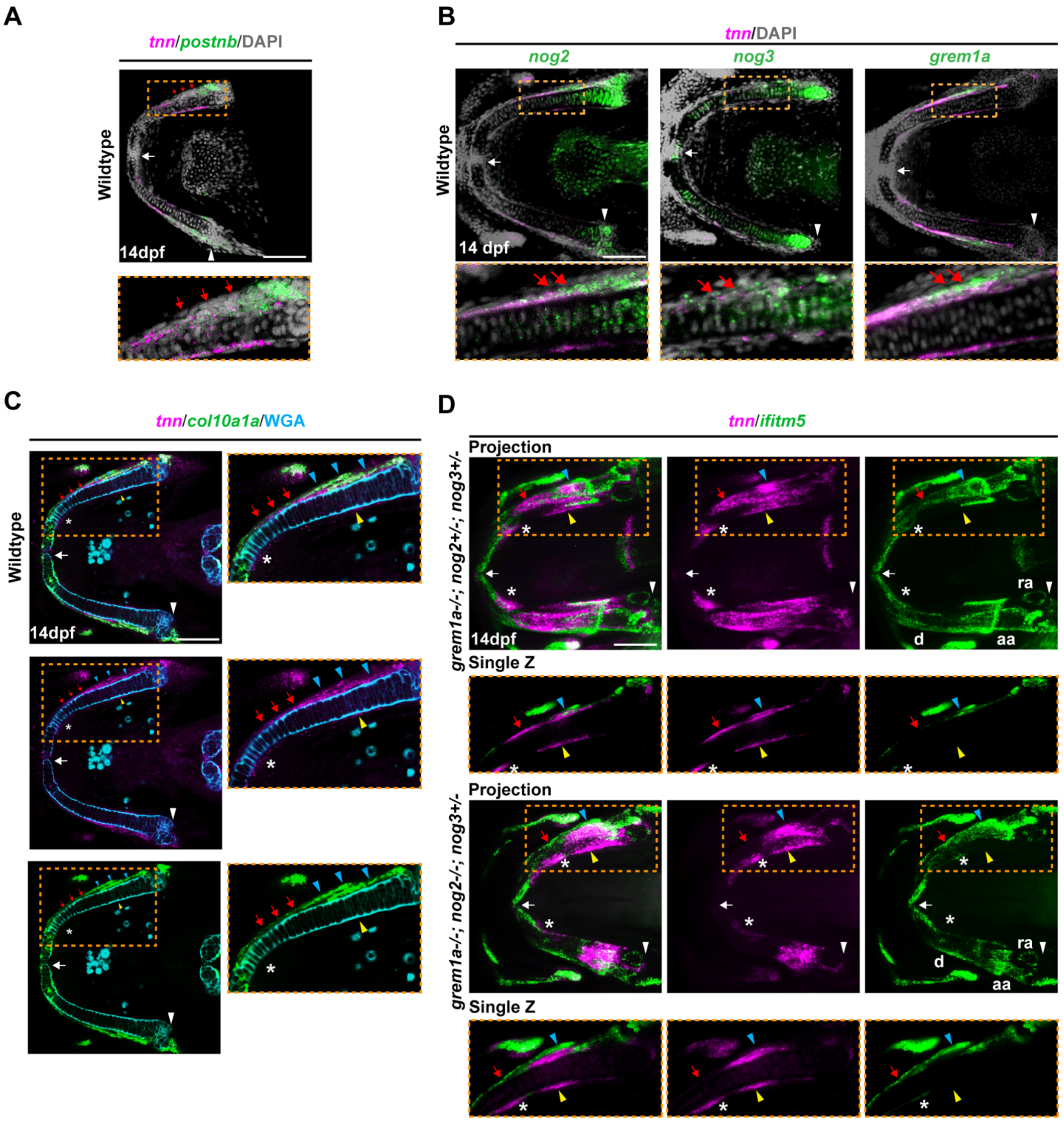


**Supp. Figure 6. Loss of BMP antagonists does not impair osteogenesis but disrupts the organization of *tnn*⁺ skeletal mesenchyme adjacent to Meckel's cartilage.** (**A**) Single Z-plane confocal images of Meckel's cartilage following in situ hybridization for *postnb* (green) and *tnn* (magenta), counter-stained with DAPI (performed in biological triplicates). Yellow dashed boxes indicate regions shown in higher-magnification insets. Red arrows denote double-positive cells. (**B**) Single Z-plane confocal images of Meckel's cartilage following in situ hybridization for *tnn* (magenta), *nog2, nog3, and grem1a* (green) and counter-stained with DAPI (gray) at 14 dpf (performed in biological triplicates). Yellow dashed boxes indicate regions shown in higher-magnification insets. Distal *tnn*+ mesenchyme co-expresses BMP antagonists (indicated by red arrows). (**C**) Single Z-plane confocal images of Meckel's cartilage following in situ hybridization for *tnn* (magenta) and *col10a1a* (green) and counter-stained with Wheat Germ Agglutinin (blue) at 14 dpf (performed in biological triplicates). Yellow dashed boxes indicate regions shown in higher-magnification insets. *tnn*+ mesenchyme extends along the Meckel's cartilage but stops before reaching *col10a1a*+ chondrocytes. Red arrows indicate the distal mesenchymal expression, blue arrowheads mark the proximal mesenchymal expression, and yellow arrowheads mark the medial mesenchymal expression. *tnn*+ tendon is labeled with an asterisk. (**D**) Maximum projection confocal images of Meckel's cartilage following in situ hybridization for *tnn* (magenta) and *ifitm5* (green) in control and dhomo; het mutant skeletons at 14 dpf (performed in biological triplicates). Yellow dashed boxes indicate regions shown in higher-magnification single Z-planes. Distal *tnn* expression adjacent to Meckel's cartilage is specifically reduced in 14 dpf dhomo; het (red arrows), whereas *tnn* expression persists in proximal mesenchyme (blue arrowheads), medial mesenchyme (yellow arrowheads), and tendon (asterisks). Dermal bones, labeled by *ifitm5* expression (green), appear comparable to controls. White arrows point to the anterior of the Meckel's cartilage and arrowheads indicated the posterior of the Meckel's cartilage. d, dentary; aa, anguloarticular; ra, retroarticular. Scale bars, 100 μm.
